# Multidimensional quantitative analysis of mRNA expression within intact vertebrate embryos

**DOI:** 10.1101/214619

**Authors:** Vikas Trivedi, Harry M.T. Choi, Scott E. Fraser, Niles A. Pierce

**Affiliations:** Division of Biology & Biological Engineering, California Institute of Technology, Pasadena, CA 91125, USA.; Translational Imaging Center, University of Southern California, Los Angeles, CA 90089, USA.; Molecular and Computational Biology, University of Southern California, Los Angeles, CA 90089, USA.; Department of Genetics, University of Cambridge, Cambridge, CB2 3EH, UK.; Division of Engineering & Applied Science, California Institute of Technology, Pasadena, CA 91125, USA.; Weatherall Institute of Molecular Medicine, University of Oxford, Oxford OX3 9DS, UK.

**Keywords:** quantitative in situ hybridization, multiplexed in situ hybridization, read-out, read-in

## Abstract

For decades, in situ hybridization methods have been essential tools for studies of vertebrate development and disease, as they enable qualitative analyses of mRNA expression in an anatomical context. Quantitative mRNA analyses typically sacrifice the anatomy, relying on embryo microdissection, dissociation, cell sorting, and/or homogenization. Here, we eliminate the tradeoff between quantitation and anatomical context, using multiplexed in situ hybridization chain reaction (HCR) to perform accurate and precise relative quantitation of mRNA expression with subcellular resolution within whole-mount vertebrate embryos. Gene expression can be queried in two directions: read-out from anatomical space to expression space reveals co-expression relationships in selected regions of the specimen; conversely, read-in from multidimensional expression space to anatomical space reveals those anatomical locations in which selected gene co-expression relationships occur. As we demonstrate by examining gene circuits underlying somitogenesis, quantitative read-out and read-in analyses provide the strengths of flow cytometry expression analyses, but by preserving subcellular anatomical context, they enable iterative bi-directional queries that open a new era for in situ hybridization.

**SUMMARY:** Multiplexed in situ hybridization chain reaction (HCR) enables quantitative multidimensional analyses of developmental gene expression with subcellular resolution in an anatomical context.

---

Traditional in situ hybridization approaches based on catalytic reporter deposition (CARD) yield high-contrast images of mRNA expression domains within whole-mount vertebrate embryos (Tautz & Pfeifle, 1989; Harland, 1991; Lehmann & Tautz, 1994; Kerstens *et al*., 1995; Nieto *et al*., 1996; Pernthaler *et al*., 2002; Kosman *et al*., 2004; Thisse *et al*., 2004; Denkers *et al*., 2004; Clay & Ramakrishnan, 2005; Barroso-Chinea *et al*., 2007; Acloque *et al*., 2008; Piette *et al*., 2008; Thisse & Thisse, 2008; Weiszmann *et al*., 2009; Ruf-Zamojski *et al*., 2015). However, the intensity of the staining is qualitative due to the nonlinear effects of CARD; furthermore, spatial resolution is often compromised by diffusion of reporter molecules prior to deposition (Tautz & Pfeifle, 1989; Thisse *et al*., 2004; Thisse & Thisse, 2008; Acloque *et al*., 2008; Piette *et al*., 2008; Weiszmann *et al*., 2009), and multiplexing is cumbersome, requiring serial staining of each target mRNA (Lehmann & Tautz, 1994; Nieto *et al*., 1996; Thisse *et al*., 2004; Denkers *et al*., 2004; Kosman *et al*., 2004; Clay & Ramakrishnan, 2005; Barroso-Chinea *et al*., 2007; Acloque *et al*., 2008; Piette *et al*., 2008). These strengths and weaknesses all derive from the enzyme-mediated deposition process responsible for signal amplification. Direct-labeled probes offer complementary tradeoffs, avoiding nonlinear signal amplification to enable quantitative, high-resolution, multiplexed studies in thin samples (Kislauskis *et al*., 1993; Femino *et al*., 1998; Levsky *et al*., 2002; Kosman *et al*., 2004; Capodieci *et al*., 2005; Chan *et al*., 2005; Raj *et al*., 2008), but often generating insufficient signal to achieve the needed contrast in thick samples such as whole-mount vertebrate embryos.

To quantify relative mRNA expression levels for defined anatomical regions within vertebrate embryos, it is necessary to destroy the sample morphology. Current approaches employ some combination of microdissection (Nawshad *et al*., 2004; Redmond *et al*., 2014; Treutlein *et al*., 2014), cell dissociation (Manoli & Driever, 2012; Jean *et al*., 2015; Petropoulos *et al*., 2016), homogenization (Axelsson *et al*., 2007; de Jong *et al*., 2010; Pena *et al*., 2014), fluorescence-activated cell sorting (Manoli & Driever, 2012; Treutlein *et al*., 2014; Allison *et al*., 2016), magnetic-activated cell sorting (Treutlein *et al*., 2014; Taylor *et al*., 2016; Allison *et al*., 2016), or lysis (Nawshad *et al*., 2004; de Jong *et al*., 2010; Redmond *et al*., 2014; Treutlein *et al*., 2014; Laranjeiro & Whitmore, 2014; Jean *et al*., 2015; Petropoulos *et al*., 2016; Allison *et al*., 2016), followed by RNA quantitation using real-time polymerase chain reaction (Nawshad *et al*., 2004; Axelsson *et al*., 2007; Pena *et al*., 2014; Laranjeiro & Whitmore, 2014; Jean *et al*., 2015), RNA sequencing (Treutlein *et al*., 2014; Petropoulos *et al*., 2016; Allison *et al*., 2016), in situ hybridization flow cytometry (Taylor *et al*., 2016; Allison *et al*., 2016), microarray hybridization (de Jong *et al*., 2010; Redmond *et al*., 2014; Jean *et al*., 2015), or hybridization barcoding (Pena *et al*., 2014; Laranjeiro & Whitmore, 2014). Owing to this fundamental tradeoff between anatomical context and quantitation, there is an unmet need for multiplexed quantitative analysis of mRNA expression with high-resolution within intact specimens.

We have shown previously that in situ hybridization chain reaction (HCR; Figure 1A) (Dirks & Pierce, 2004; Choi *et al*., 2010) enables straightforward multiplexing, high contrast, and subcellular resolution when mapping target mRNAs within complex specimens (Choi *et al*., 2014; Choi *et al*., 2016). In situ HCR uses DNA probes complementary to mRNA targets to trigger the self-assembly of fluorophore-labeled DNA HCR hairpins into tethered fluorescent amplification polymers. Using a library of orthogonal HCR amplifiers, signal amplification is performed for all targets simultaneously. Here, we demonstrate the crucial property that the amplified HCR signal is proportional to the number of target mRNAs per subcellular imaging voxel (Figure 1B), enabling accurate and precise relative quantitation within intact vertebrate embryos.

**Fig. 1:**
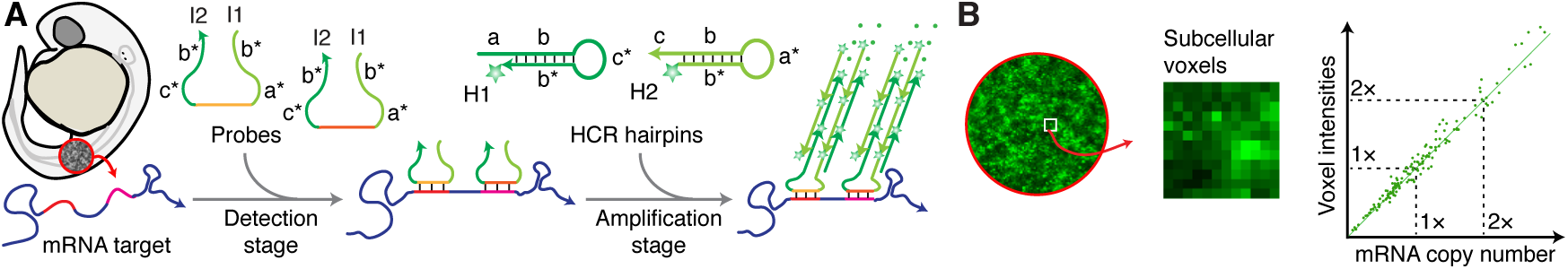
Quantitative in situ hybridization chain reaction (HCR). (A) Two-stage protocol independent of the number of target mRNA species (Choi *et al*., 2014; Choi *et al*., 2016). Detection stage: DNA probes carrying DNA HCR initiators (I1 and I2) hybridize to target mRNAs and unused probes are washed from the sample. Amplification stage: metastable fluorophore-labeled DNA HCR hairpins (H1 and H2; green stars denote fluorophores) penetrate the sample without interacting; initiators trigger chain reactions in which H1 and H2 hairpins sequentially nucleate and open to assemble into tethered fluorescent amplification polymers; unused hairpins are washed from the sample. (B) Conceptual schematic: for subcellular voxels within whole-mount vertebrate embryos, HCR signal scales approximately linearly with mRNA abundance, enabling quantitative analysis of mRNA expression in an anatomical context.

We demonstrate in situ mRNA quantitation by examining somitogenesis in the zebrafish embryo. In all vertebrates, somites provide one of the first outward appearances of the metameric body plan, periodically pinching off from the presomitic mesoderm (PSM) in bilaterally symmetrical pairs as precursors to the axial muscles and vertebral column (Oates *et al*., 2012). To date, detailed studies of the gene dynamics underlying somitogenesis have relied heavily on examination of 1-channel (Oates & Ho, 2002; Henry *et al*., 2002; Mara *et al*., 2007; Gomez *et al*., 2008; Ferjentsik *et al*., 2009; Choorapoikayil *et al*., 2012; Schroter *et al*., 2012) and 2-channel (Holley *et al*., 2000; Oates & Ho, 2002; Jülich *et al*., 2005) CARD images that display expression of 1 or 2 mRNAs per embryo. Here, using in situ HCR, we demonstrate quantitative subcellular analyses of 4 mRNAs in the same embryo. Read-out analyses reveal quantitative changes in gene co-expression ratios as somites mature moving away from the PSM, and read-in analyses reveal the anatomical locations where distinct gene co-expression relationships occur.

## RESULTS

### Accuracy and precision assessed by redundant detection

Subcellular mRNA quantitation within thick samples such as whole-mount vertebrate embryos has not been rigorously verified for any method. We chose to meet this challenge by exploiting the ease of multiplexing using in situ HCR: we redundantly detect the same target mRNA using two distinct probe sets, each of which carries initiators for orthogonal HCR amplifiers labeled with spectrally distinct fluorophores (Figure 2A). This experimental design provides an avenue for validating that HCR signal scales linearly with the number of target mRNAs per voxel, without requiring knowledge of the absolute number of targets in any voxel.

**Fig. 2:**
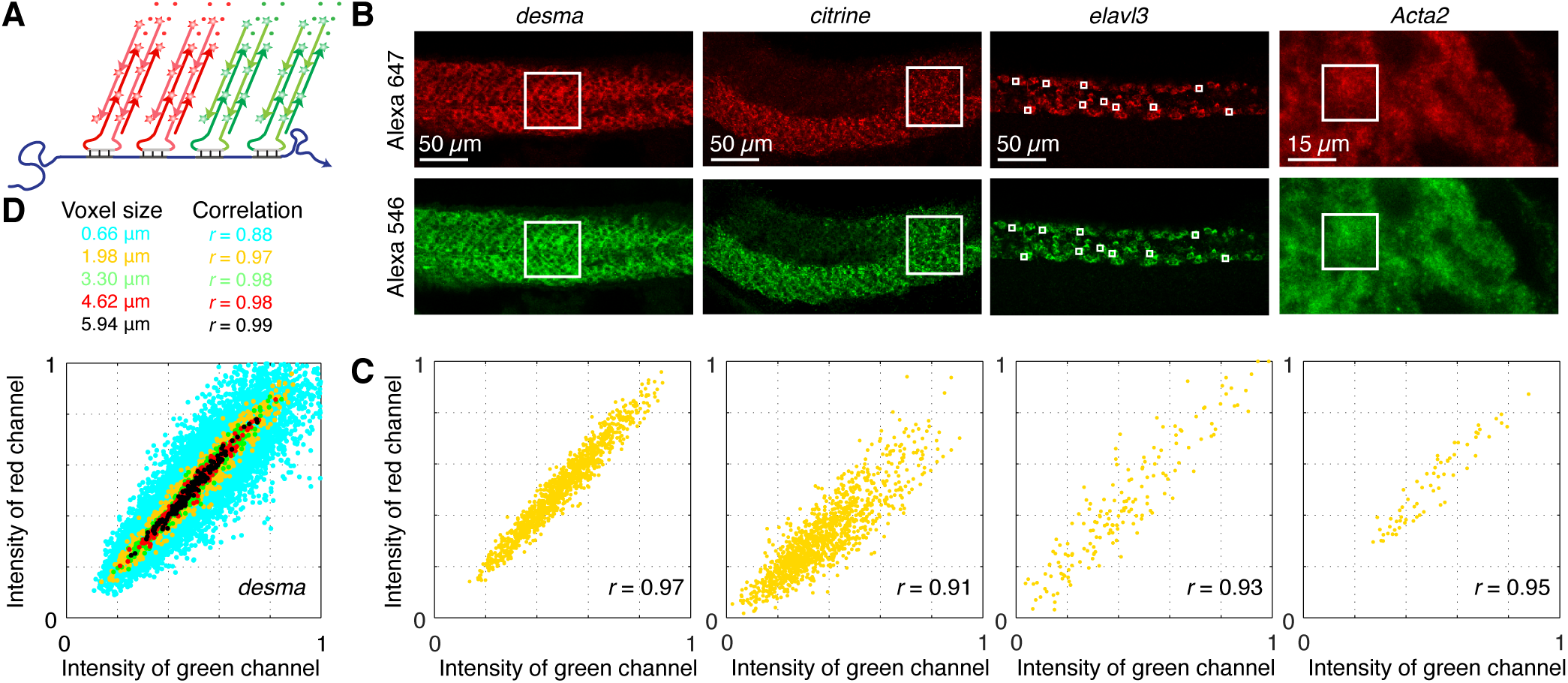
Accuracy and precision assessed by redundant detection. (A) Each target mRNA is detected using two probe sets, each initiating an orthogonal and spectrally distinct HCR amplifier (red channel: Alexa 647, green channel: Alexa 546). (B) Two-channel redundant detection of four target mRNAs: *desma*, *Gt(desma-citrine)*, and *elavl3* in whole-mount zebrafish embryos (fixed 26 hpf) and *Acta2* in a whole-mount mouse embryo (fixed E9.5). Confocal microscopy: 0.7 × 0.7 *µ*m pixels (*desma, citrine, elavl3*) or 0.07 × 0.07 *µ*m pixels (*Acta2*). (C) Highly correlated normalized signal (Pearson correlation coefficient, r) for 2 × 2 × 2 *µ*m voxels in the selected regions of panel B. Accuracy: linear distribution with zero intercept. Precision: scatter around the line. (D) Scatter as a function of voxel size for desma. See Supplementary Section S2.2 for a discussion of the effect of averaging on quantitative precision and Supplementary Section S2.3 for additional data.

To assist with our interpretation of this 2-channel test, let *ni* denote the (unknown) number of target molecules in voxel *i* and let *xi* and *yi* denote the normalized HCR signal in voxel *i* falling in the interval [0, 1] for each of the two channels. Suppose *x*_*i*_ and *y*_*i*_ are each proportional to *n*_*i*_:

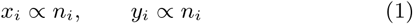

In this ideal scenario, a scatter plot of (*xi, yi*) pairs would fall exactly on a line with intercept zero, and relative quantitation of a target mRNA for any pair of voxels *j* and *i* could be calculated exactly as the ratio of voxel intensities in either channel:

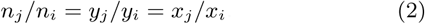

In practice, when the signal approximately satisfies (1), the (*xi, yi*) pairs will be scattered around a line with intercept zero; the accuracy of relative quantitation using (2) will then depend on the deviation of the underlying relationship from ideality (linear with zero intercept), while the precision of (2) will depend on the scatter around the line.

Using this approach, we perform 2-channel redundant detection of four different target mRNAs within whole-mount zebrafish or mouse embryos (Figure 2B), observing highly correlated subcellular voxel intensities (Figure 2C; Pearson correlation coefficient 0.91 ≤*r* 0.97 for 2*×*2*×*2 *µ*m voxels). For each target mRNA, variation along the diagonal indicates biological variation in expression levels between voxels. Despite numerous potential sources for non-ideality, the accuracy is high (clear linear relationship with intercept near zero) and the precision is very good (scatter of approximately 5% to 25% of the dynamic range depending on the target mRNA). The scatter in each channel arises from non-uniformity in the background per voxel as well as from non-uniformity in the signal generated per target molecule (resulting from variation in probe hybridization yields (Raj *et al*., 2008; Shah *et al*., 2016) and HCR amplification polymer lengths (Dirks & Pierce, 2004; Choi *et al*., 2010; Choi *et al*., 2014)). Scatter is reduced by averaging neighboring pixel intensities, enabling dramatic improvements in precision while still retaining subcellular resolution (Figure 2D). This averaging effect is also evident in the high precision achieved using HCR to perform relative and absolute quantitation of miRNAs in northern blots (Schwarzkopf & Pierce, 2016), where the size of the voxel is effectively the size of the band being quantified. We expand upon the origins and properties of this important averaging effect in Supplementary Section S2.2.

### Measuring a 2-fold difference in mRNA levels

We next tested our ability to use in situ HCR to discriminate a known difference in mRNA levels between embryos. The FlipTrap line *Gt(desma-citrine)* (Trinh *et al*., 2011) provides an ideal test setting as the expression level for citrine is expected to be approximately 2-fold higher in homozygous vs heterozygous embryos (expressed from 2 or 1 transgenic alleles), while desma expression is expected to be similar in both genotypes (expressed from both alleles regardless). Extending our 2-channel redundant detection approach, we detect citrine with one probe set and desma with a second probe set (Figure 3A). Comparing mean citrine expression levels within regions of high expression (Figure 3B) yields a homo/hetero expression ratio of 2.0 *±* 0.5 (Figure 3C). This roughly 2-fold difference in expression level is also evident in the distinct slopes observed in scatter plots of subcellular voxel intensities (Figure 3D). The ability to discriminate 2-fold changes in mRNA expression in an anatomical context is important, for example, in evaluating perturbations or candidate drugs intended to alter gene expression.

**Fig. 3:**
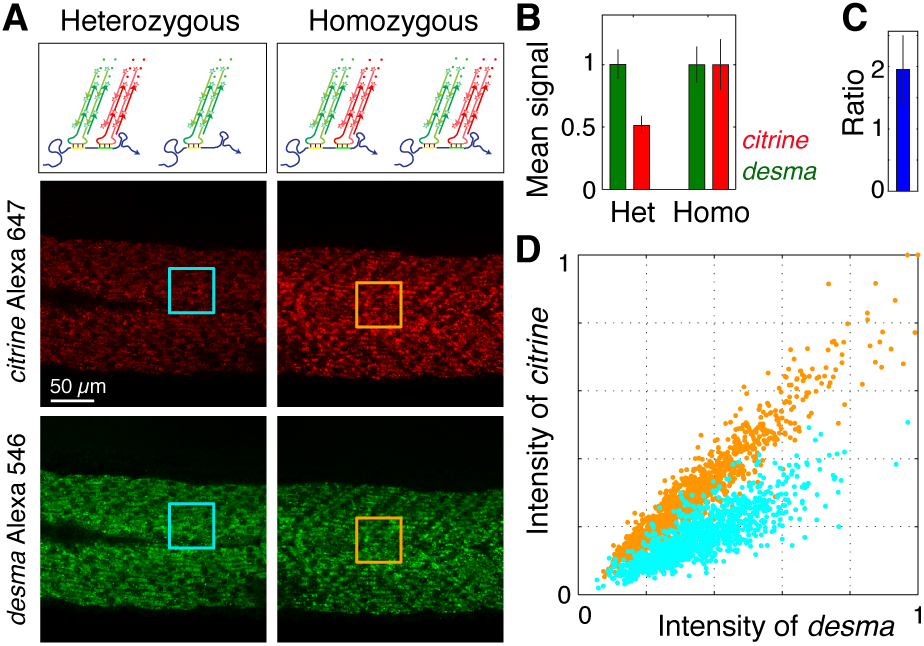
Measuring a 2-fold difference in mRNA levels. (A) Two-channel imaging of citrine (red channel: Alexa 647) and desma (green channel: Alexa 546) target mRNAs in homozygous *Gt(desma-citrine)*^*ct122a/ct122a*^ embryos and heterozygous *Gt(desma-citrine)*^*ct122a/+*^ embryos. Confocal microscopy: 0.7 × 0.7 *µ*m pixels. Whole-mount zebrafish embryos fixed 26 hpf. (B) Normalized signal for citrine (red) and desma (green) targets in depicted regions of homozygous and heterozygous embryos (mean standard deviation, N=3 embryos). (C) Ratio of citrine target in homozygous vs heterozygous embryos (mean standard deviation, N=3 embryos). (D) Normalized signal for 2 × 2 × 2 *µ*m voxels within the selected regions of panel A. See Supplementary Section S2.4 for additional data.

### Quantitative read-out from selected anatomical locations to multidimensional expression space

We next apply in situ HCR to multidimensional analyses of zebrafish somitogenesis to explore the power of mRNA quantitation in an anatomical context. Figure 4A displays a 4-channel image for four target mRNAs expressed as the somites emerge and are displaced from the PSM: two cyclic segmentation genes (*her7* and *her1*) and two muscle genes (*myod1* and *tpm3*). Figure 4B presents expression profiles in a strip spanning five regions of interest (somites S7, S8, S9, S10, and the PSM), revealing the relative expression levels from the older somite (S7) to the tissue soon to become a somite: approximately (100%, 75%, 50%, 25%, 0%) for *myod1* (magenta curve) and approximately (0%, 0%, 0%, 50% 100%) for *her7* (red curve). These profiles provide a first quantitative glimpse of the strongly anti-correlated bulk expression trends for *myod1* and *her7*, but do not take advantage of the subcellular resolution of the data.

**Fig. 4:**
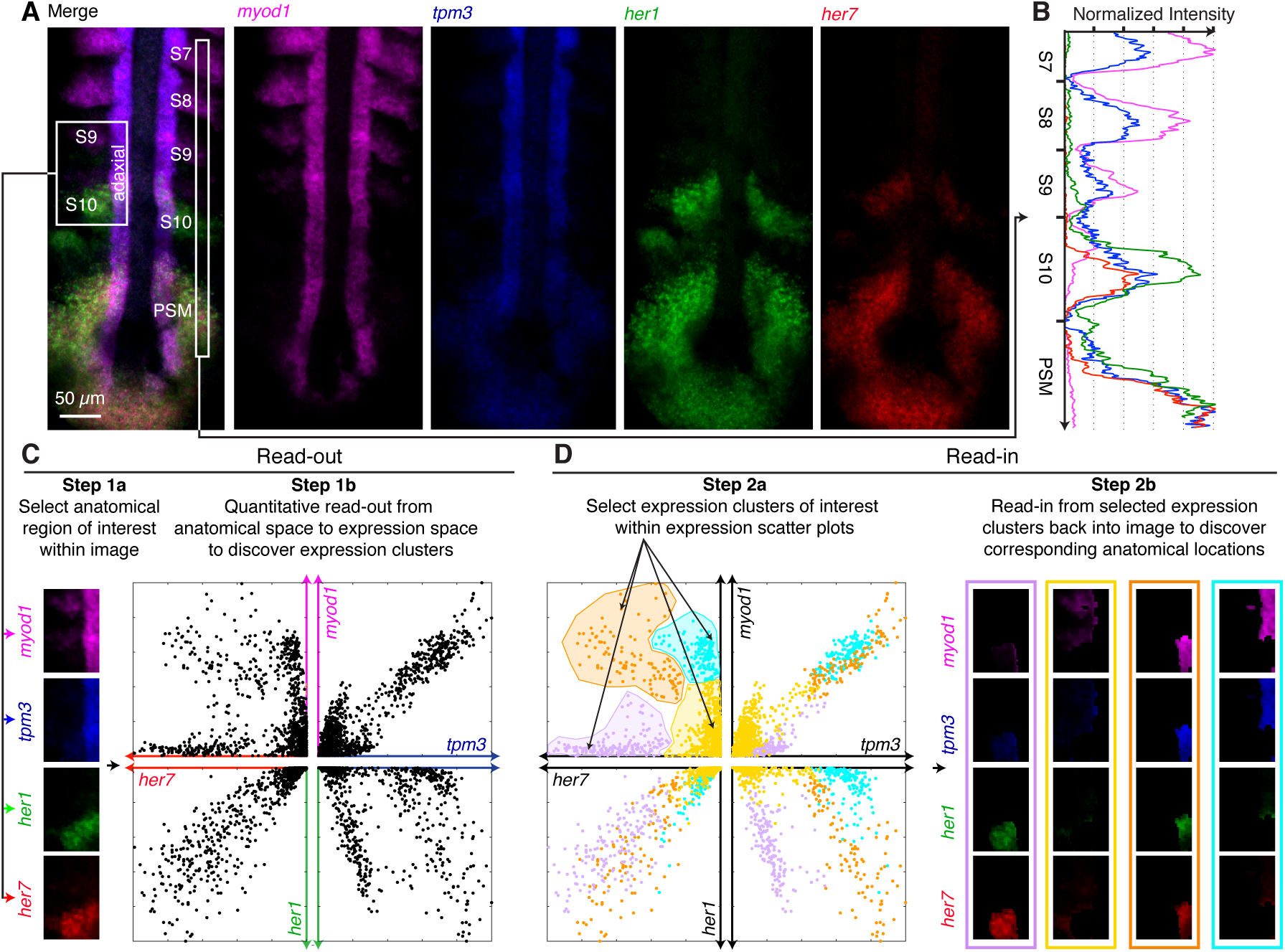
Quantitative read-out and read-in. (A) Four-channel quantitative image for four target mRNAs in a whole-mount zebrafish embryo. Confocal microscopy: 0.7 × 0.7 *µ*m pixels, mean intensity over 5 focal planes. Embryo fixed 10 hpf. (B) Normalized expression profiles for four target mRNAs along a strip of interest (see panel A) crossing four somites (S7, S8, S9, S10) and the presomitic mesoderm (PSM). (C) Read-out from a region of interest (see panel A) within a 4-channel image (left) to pairwise expression scatter plots (right), revealing distinct expression clusters with different expression characteristics. Each point within an expression scatter plot represents normalized voxel intensities for a pair of target mRNAs. Voxel size: 2 × 2 × 6 *µ*m. (D) Read-in from pairwise expression scatter plots (left) to a 4-channel image (right), revealing the anatomical locations corresponding to 4 expression clusters of interest. Expression clusters selected in the *her7-myod1* quadrant; cluster shading (yellow, purple, orange, blue) propagated to other three quadrants. See Supplementary Section S2.5 for additional data.

To perform subcellular quantitative analyses, we display scatter plots of voxel intensities reminiscent of multiplexed fluorescence-activated cell sorting (FACS) data. Figure 4C demonstrates read-out from a region of interest in the 4-channel image (in this case, a rectangle containing somites S9, S10, and a longitudinal stripe of adaxial cells) to scatter plots of normalized voxel intensities for pairs of target mRNAs. Strikingly, these expression scatter plots reveal well-defined expression clusters with differing slopes and amplitudes corresponding to voxels with related expression characteristics. For example, the *her7* vs *myod1* quadrant reveals one cluster with low *myod1* and variable *her7* levels, a second cluster with low *her7* and variable *myod1* levels, and a third cluster with correlated variable expression of both targets.

### Quantitative read-in from selected expression clusters to anatomical locations

Unlike FACS analysis where the display of expression clusters would be the final product, in situ HCR maintains the anatomical context, permitting us to map expression clusters of interest back into the embryo to interactively investigate their physical positions. Figure 4D demonstrates read-in from four selected expression clusters in the *her7* vs *myod1* quadrant (shaded purple, yellow, orange, and blue) back into the embryo, segmenting the image into four expression domains. The purple expression cluster maps to the posterior portion of S10, and the yellow expression cluster maps to S9 and the anterior portion of S10, placing the boundary between these expression domains in the middle of the youngest somite. The orange and blue expression clusters map to the adjacent portions of the longitudinal stripe of adaxial cells. Cluster shading can be propagated between the quadrants in the expression scatter plots, which is akin to projecting the voxel intensities onto 4 axes, but without the difficulty of 4-dimensional visualization. For example, the well-separated purple and yellow clusters identified in the *her7* vs *myod1* quadrant partition the complex cluster near the origin in the *myod1* vs *tpm3* quadrant into two clusters with differing slopes (Figure 4D). Similar read-in analyses can be performed for expression clusters identified in any quadrant or combination of quadrants, permitting a detailed examination of the spatial organization of distinct genetic circuit states (Supplementary Figures S29-S35).

### Quantitative snapshots of gene co-expression changes during somite formation and maturation

Somites are rhythmically and sequentially generated by the presomitic mesoderm (PSM), resulting in a developmental time course being reflected within a single embryo (somite S7 is developmentally more mature than S8, which is more mature than S9, and so on), with new somites emerging at approximately 30-minute intervals in zebrafish (Oates *et al*., 2012). Having previously used expression clusters within the scatter plots to identify expression regions within the image (Figure 4D), we now reverse the direction of information flow and use each somite within the image to identify expression clusters within the scatter plots. This approach yields quantitative snapshots of gene coexpression changes as the somites form and mature. Figure 5A depicts 4-channel read-out from five regions (S7, S8, S9, S10, PSM) to expression clusters shaded by their anatomical location (black, blue, green, yellow, and red). The *myod1* vs *tpm3* quadrant reveals striking changes in slope and amplitude during this maturation process (Figure 5B): a low slope in the PSM jumps to a higher slope in maturing somites S9, S8, and S7; within these three older somites, amplitude increases monotonically with somite maturity. The youngest somite (S10) exhibits expression clusters with both slopes, consistent with the observations of Figure 4D. The similar slopes and small intercept for the three older somites indicate that the ratio of *myod1* to *tpm3* expression remains approximately constant as the somites mature. Interestingly, replicate embryos of nominally the same age capture slightly different developmental stages within the oscillatory somitogenesis circuitry, revealing expression clusters with slopes and amplitudes reflecting related but different circuit states (Supplementary Figure S37). Within each embryo, left and right somites are similar, serving as technical replicates for slope and amplitude measurements (Supplementary Figures S37-S41).

**Fig. 5:**
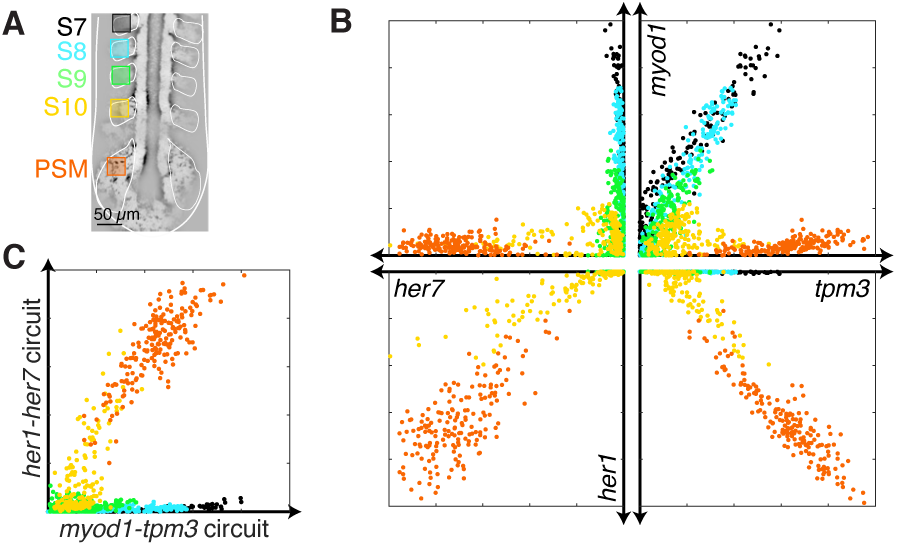
Quantitative snapshots of gene co-expression changes during somite formation and maturation. (A) Anatomical regions of interest within somites S7, S8, S9, S10 and the presomitic mesoderm (PSM). (B) Expression scatter plots for four target mRNAs shaded by anatomical regions of panel A. (C) Subcircuit expression scatter plots. Amplitude of *her1-her7* subcircuit 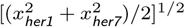 vs amplitude of *myod1-tpm3* 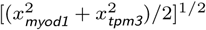 for the anatomical regions of panel A. *x* denotes normalized signal for each target mRNA. Confocal microscopy: mean intensity over 5 focal planes, 2 × 2 × 6 *µ*m voxels. Embryo fixed 10 hpf. See Supplementary Section S2.6 for additional data.

Discerning two subcircuits emerging in the expression clusters of Figure 5B, we project the voxel intensities onto new axes to examine somitogenesis through the lens of a *her1-her7* subcircuit and a *myod1-tpm3* subcircuit (Figure 5C). There is a crossover within youngest somite (S10) between dominance of the *her1-her7* subcircuit in the PSM and the *myod1-tpm3* subcircuit in maturing somites S9, S8, and S7, leading to nearly orthogonal expression clusters in Figure 5C. Subcircuit projections can be performed onto axes representing an arbitrary number of circuit elements, compactly summarizing a large quantity of high-dimensional expression information.

## DISCUSSION

In situ HCR dramatically expands the capabilities of in situ hybridization as a research tool. mRNA expression levels can be compared within an anatomical region (Figure 2), between regions (Figure 4B), and between embryos (Figure 3). Quantitative discovery is enabled in two directions: read-out from multi-channel images to co-expression scatter plots reveals well-defined expression clusters with differing slopes and amplitudes (Figure 4C); read-in from expression scatter plots to multi-channel images reveals the anatomical locations in which select co-expression relationships occur (Figure 4D). In one direction, anatomical locations within the image can be used to identify expression clusters within scatter plots (Figure 5), and in the other direction, expression clusters within the scatter plots can be used to identify anatomical locations within the image (Figure 4D). These capabilities follow from two crucial properties of in situ HCR: multiplexing spreads the voxel data out onto multiple expression axes to enable identification of well-segregated expression clusters; quantitation generates expression clusters that display slopes and amplitudes revealing similarities and differences between genetic circuit states. Projection of expression scatter plots onto subcircuit axes representing one or more target mRNAs facilitates quantitative dissection of the underlying regulatory circuitry across multiple anatomical regions. Collectively, these capabilities open a new era for in situ hybridization, enabling quantitative bidirectional interrogation of anatomical locations and expression clusters in the study of vertebrate development and disease.

## METHODS

### Performing read-out/read-in analyses

See Figure 6 for an illustration of the read-out/read-in workflow:

**Fig. 6:**
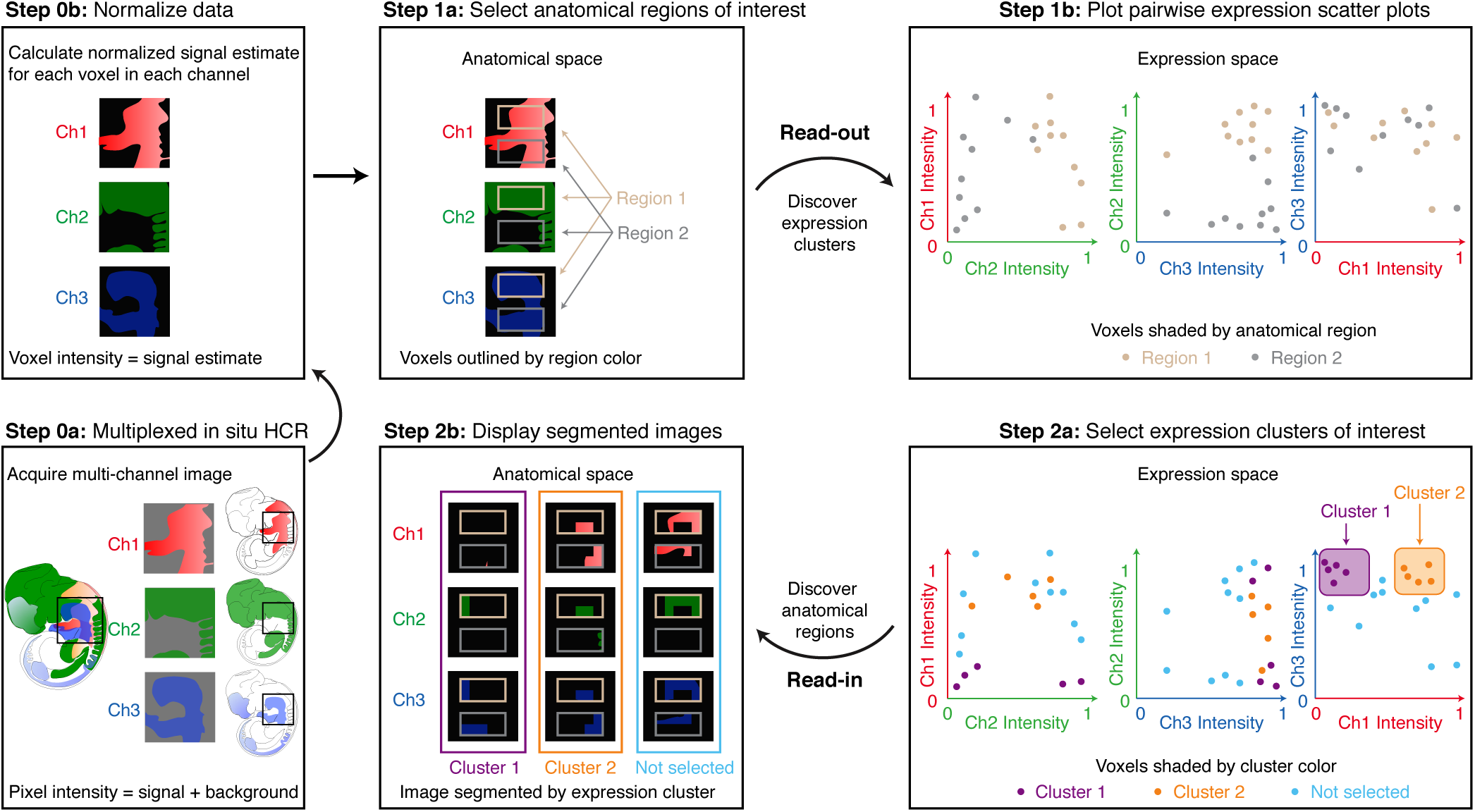
Work flow for quantitative read-out and read-in analyses using in situ HCR. Step 0: Acquire and normalize data. Step 1: Read-out from anatomical space to expression space. Step 2: Read-in from expression space to anatomical space. If desired, Steps 1 and 2 can be performed iteratively, moving back and forth between regions of interest in anatomical space and clusters of interest in expression space.

**Step 0:** Acquire and normalize data. a) Perform a multiplexed in situ HCR experiment. Image the expression patterns for *N* target mRNAs using an *N*-channel experiment in a single specimen (Choi *et al*., 2016) (3 channels depicted in Figure 6). b) Calculate normalized voxel intensities for each channel. Average pixel intensities to create raw voxel intensities for each channel (Section 1.4.1) and then calculate the normalized signal estimate for each voxel in each channel (Section S1.4.3). The same normalization should be applied to all images to enable quantitative comparison between embryos.

**Step 1:** Read-out from anatomical space to expression space. a) Select anatomical regions of interest within a multi-channel image (2 regions of interest depicted in Figure 6: tan and gray rectangles). b) Plot expression scatter plot for each pair of channels: for each voxel in the regions of interest, plot the normalized signal intensity shaded by region color (tan or gray dots in Figure 6; see also examples in Figure 4C and 5B).

**Step 2:** Read-in from expression space to anatomical space. a) Select expression clusters of interest (magenta and orange clusters selected in Ch3 vs Ch1 scatter plot, dots not selected shaded blue). b) Redisplay the multichannel image showing only the voxels in one expression cluster at a time (images bounded by magenta, orange, or blue rectangles; see also example of Figure 4D).

## Acknowledgments

This work is dedicated to our friend and colleague, Robert M. Dirks, who tragically lost his life in the Metro-North train crash in New York in 2015. We thank L.A. Trinh and B.R. Wolfe for helpful discussions and L.A. Fletcher, C. Paquette, and D. Mayorga for fish care.

## Author contributions

Methodology: V.T., H.M.T.C., S.E.F., N.A.P.; Software: V.T.; Validation: V.T.; Investigation: V.T., H.M.T.C.; Writing - original draft: V.T., N.A.P.; Writing - review and editing: V.T., H.M.T.C., S.E.F., N.A.P.; Visualization: V.T., S.E.F., N.A.P.; Supervision: S.E.F., N.A.P.; Project administration: N.A.P.; Funding acquisition: S.E.F., N.A.P.

## Supplementary information

Supplementary information available.

